# Determinants of Nutritional Status among Children Under five age in Ethiopia: A Further Analysis of the Ethiopian Demographic and Health Survey (EDHS) 2016 Data

**DOI:** 10.1101/698308

**Authors:** Zerihun Yohannes Amare, Mossa Endris Ahmed, AdeyBelete Mehari

**Author notes:** ZYM contributed mainly to this study. MEH and ABM contributed equally to this work. Corresponding author; /.

## Abstract

Child malnutrition is an underlying cause for almost half (45%) of child deaths, particularly in low socioeconomic communities of developing countries like Ethiopia. Globally, in 2018, 149 million children under five were stunted and over 49 million children were wasted. In Ethiopia, from the year 2005 to 2016, there is a decrease in stunting from 47% to 39%, but the prevalence of wasting changed little over the same time period (11% to 10%). Despite efforts made by the Ethiopian government and improvements in reducing malnutrition, the current rate of progress is not fast enough to have reached the global target by 2025.The aim of this study was to examine the determinants of nutritional status among children under five in Ethiopia. This study used data from 2016 Ethiopia Demographic and Heath Survey (EDHS) to examine determinants of nutritional status among children under five (0-59 months). This study used stunting and wasting as dependent variables for the analysis. Children’s, mothers, households, and environmental characteristics were used as determinant variables. Children not alive, and other missing values were considered as missing and was not included in the analyses. Sample weights were applied in all analysis due to the two stage cluster sampling design in the EDHS datasets. Multicollinearity among independent variables were checked. Logistic regression was used to analyse the determinants of nutritional status among under five age children. Bivariate analysis was also used to analyse the association between the dependent and independent variables. The chi-square test used to see the significance of association. The level of significance for the analysis was p<0.05. Age, and sex of child, educational status and body mass index, and short stature of mothers, residence, region, wealth quintile, toilet facilities and fuel types of households’ have significant association with stunting and wasting. However, mother’s short stature has significant association with only stunting. The study found child, maternal, household and environmental characteristics were significantly associated with stunting and wasting among of children under five. This implies a multi-sectorial and multidimensional approach is important to address malnutrition in Ethiopia.

## INTRODUCTION

Malnutrition during childhood is the outcome of insufficient food intake, diarrhea and other infections, lack of sanitation, low parental education (1, 2). The poor diets and disease are due to food insecurity, inadequate maternal and child care, poor health services and environment(3). It causes measurable adverse effects on body function and clinical outcome(4). This problem leads to most of the anthropometric deficits to happen among under five children in the world’s least developed countries(5). Despite existing interventions to address child malnutrition, it is still a major global public health problem (6). Child malnutrition is an underlying cause for almost half (45%) of child deaths, particularly in low socioeconomic communities of developing countries(7). Globally, in 2018, 149 million children under five suffer were stunted and over 49 million children under five were wasted (8).

In sub Saharan Africa the prevalence of stunting is declining but is still over 30% (8). Several African countries have updated their national nutrition policies, strategies and action plans(9) in order to address malnutrition. Among sub-Saharan African countries, the prevalence of wasting and stunting in Ethiopia is 10% and 38% respectively (10). Within Ethiopia there is a regional variation in terms of stunting and wasting. Amhara, Benishangul-Gumuz, Afar and Dire Dawa are the most affected by child stunting (41-46%), whereas wasting is highest in Somali (23%), Afar (18%), and Gambella (14%)(10). The current rate of progress is not fast enough to have reached the global target by 2025. Thus, to achieve the global malnutrition target for 2025, a situational analysis to determine how many children under age five years are stunted and wasted and assess key determinants of malnutrition in specific social and geographical locations are required (11). This type of analysis will be an evidence for program intervention so that programmic actions are tailored to address the contextual needs in Ethiopia.

At the policy and program level, Ethiopia has many strategies and programs to reduce malnutrition as part of its national development agenda. Some of the major strategies and programs include: the growth and transformation plan (GTP), National Nutrition Plan (NNP), the Seqota Declaration (SD), National Food Security Strategy, Nutrition Sensitive agriculture (NSa) strategy, school health and Nutrition Strategy (ShNS), the Productive Safety Net Program (PSNP), and Food Safety and Quality related regulatory activities (12). The Seqota Declaration (SD) adopted the Sustainable Development goal 2 (SDG2) with the aims to end hunger achieve food security and improve nutrition, and promote sustainable agriculture by 2030 (13).

The Government of Ethiopia through Ministry of health also launched the health extension program in 2003 to achieve the country’s progress in meeting millennium development goal. The health extension workers are mainly to improve access to care in rural communities. The health extension workers spend 15% of their time with infants and children under age five (14). All of these listed efforts have brought positive impact in improving food and nutrition security (12). However, there was a fragmented approach leading to inadequate multi-sectorial coordination in responding to food and nutrition demand. As a result, severe malnutrition have remained to be a serious challenge (12). Therefore, the rationale of the food and nutrition policy is to use fragmented efforts and resources in an integrated and clear directions and strategies so as to bring better results. Thus, this finding will be used as source of information to implement the newly introduced national food and nutrition policy.

## Materials and Methods

### DATA

This study used the 2016 EDHS dataset, which was conducted by the Central Statistical Agency (CSA), of Ethiopia, with technical support from ICF. The DHS is a population-based household survey designed to provide representative data for the country as a whole and for nine regional states and two city administrations of Ethiopia. Finally, the number of live children 0 to 59 months, with anthropometry data were considered to analyse determinants of nutritional status among under five children in Ethiopia.

### Variables

#### Dependent variables

The three anthropometric indicators mostly used for monitoring malnutrition in children are: stunting (low height-for-age), underweight (low weight-for-age); and wasting (low weight-for height). However, underweight is the composite of stunting and wasting, so this study used stunting and wasting for the analysis. They were defined using the WHO (2006) child growth standards. Stunting and, wasting were coded as binary variables based on the standard definitions.

#### Independent Variables

As it is illustrated in the conceptual framework (Figure1), to analyse the determinants of nutritional status among children under five the study used; 1) Child characteristics such as age (less than 6 months, 6-11 months, 12-17 months, 18-23 months, and 24-59 months), sex (male/female), birth order (first, second-fourth, five and greater) and perceived birth weight (very small, smaller than average, average, and very large or larger than average). 2) Mother’s characteristics such as marital status (never married, married, widowed/divorced/separated), age at current birth (12 to19, 20 to 49), educational status (no education, primary, secondary and above), working status (yes, no), body mass index (underweight, normal, overweight), and maternal stature (normal stature, short stature). Due to multicoliniarity problem preceding birth interval was removed from the analysis. 3) Household characteristics like place of residence (urban/rural), region (nine regions and two city administrations), wealth quintile (poorest, poorer, middle, rich, and richest), water sources (improved, non-improved sources), toilet facilities (improved, non-improved toilet) and cooking fuel types (traditional, modern fuel) were considered as independent variables.

Wealth quintiles in the DHS were calculated based on consumer goods such as television, bicycle or car. Household characteristics such as toilet facilities, source of drinking water, and flooring materials were also considered in calculating the wealth index. These scores were derived using principal component analysis. Based on the EDHS 2016 report, the water sources and toilet facilities were recoded as improved and non-improved water sources and toilet facilities. Piped water, public taps, standpipes, tube wells, boreholes, protected dug wells and springs, and rainwater were considered as improved water. Non-shared toilet of the following types: flush/pour flush toilets to piped sewer systems, septic tanks, and pit latrines; ventilated improved pit (VIP) latrines; pit latrines with slabs; and composting toilets are improved toilet facilities and considered for this analysis. Household fuel type was also categorized as modern and traditional fuel. Electricity, natural gas, biogas, and kerosene was categorized as modern fuel. Whereas, charcoal, wood, animal dung, and other agricultural crops and straw were also considered as traditional fuel.

#### Statistical analysis

Descriptive statistics were used to describe and illustrate the background characteristics of children 0 to 59 months. Stata 15.1 version was used to analyse the data. Sample weights were applied in all analysis due to the two stage cluster sampling design in the EDHS dataset. Multicollinearity among independent variables were checked using variance inflation factors (VIF). Multiple logistic regression was used to determine the associations between predictors and nutritional outcomes after adjusting for other covariates. The study also used Pearson chi-square test for bivariate analysis.

## RESULTS

For this analysis, the study used the 2016 DHS dataset. The missing values were not included in the analyses. Not alive children (635), with no anthropometry data from stunting condition (1000), and wasting condition (927) children were also omitted from the analysis.

Table 1 shows, the majority of mothers were married (95%), age group 20-49 (90 %) and non-working mothers (72 %). The majority (74%) of mothers are normal in their body mass index (BMI). The majority (61%) of children were age 24 to 59 months, more than half (51%) were male in their sex and only 18% were first birth children. The majority (86%) of households were male headed and 89% were reside in the rural areas of Ethiopia. The majority (83 %) of children were located only in three regions; 19% from Amhara, 44% from Oromia, and 21% from SNNP. Nearly half (46 %) of children were from poor households.

**Table 1.**
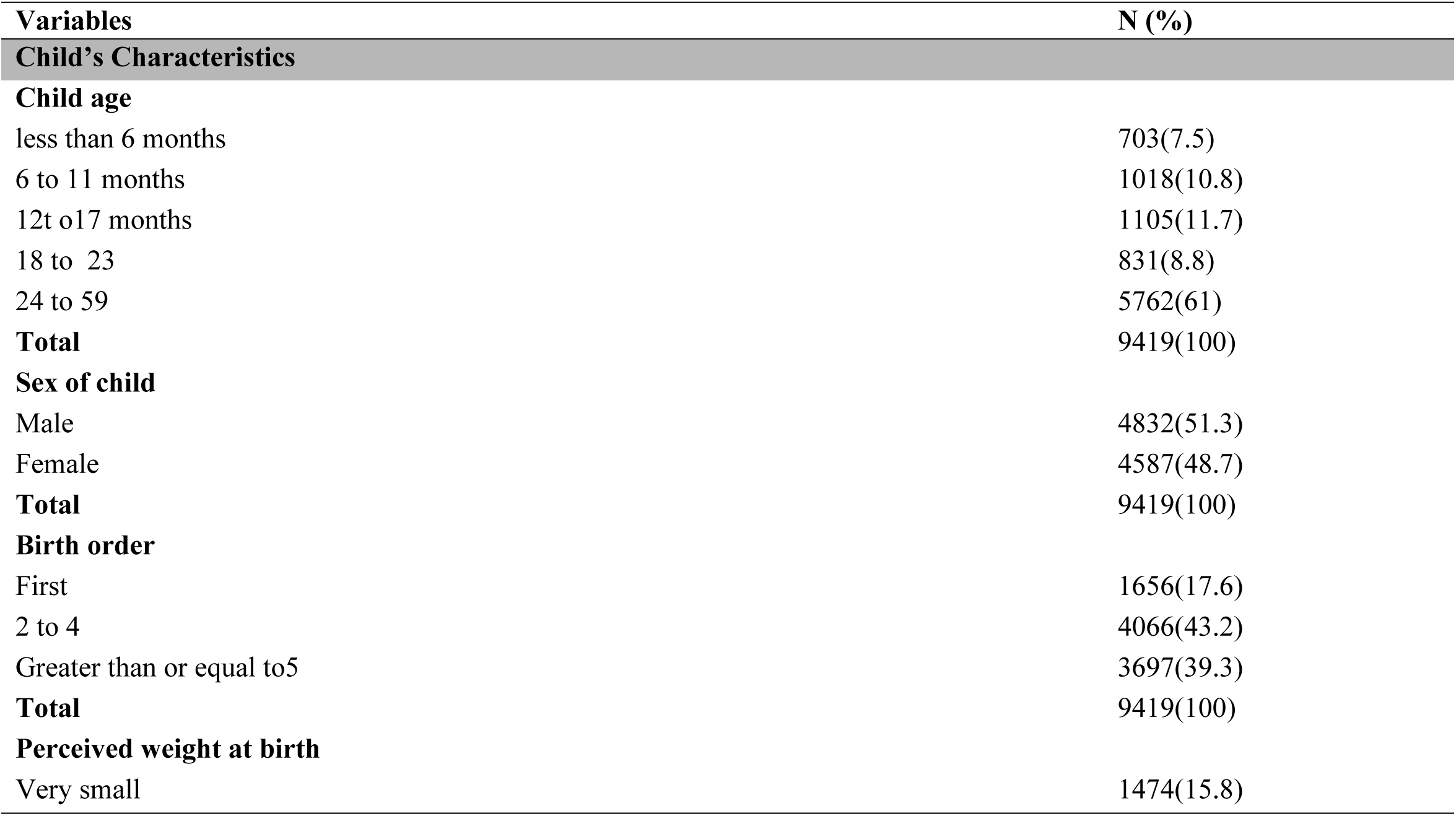
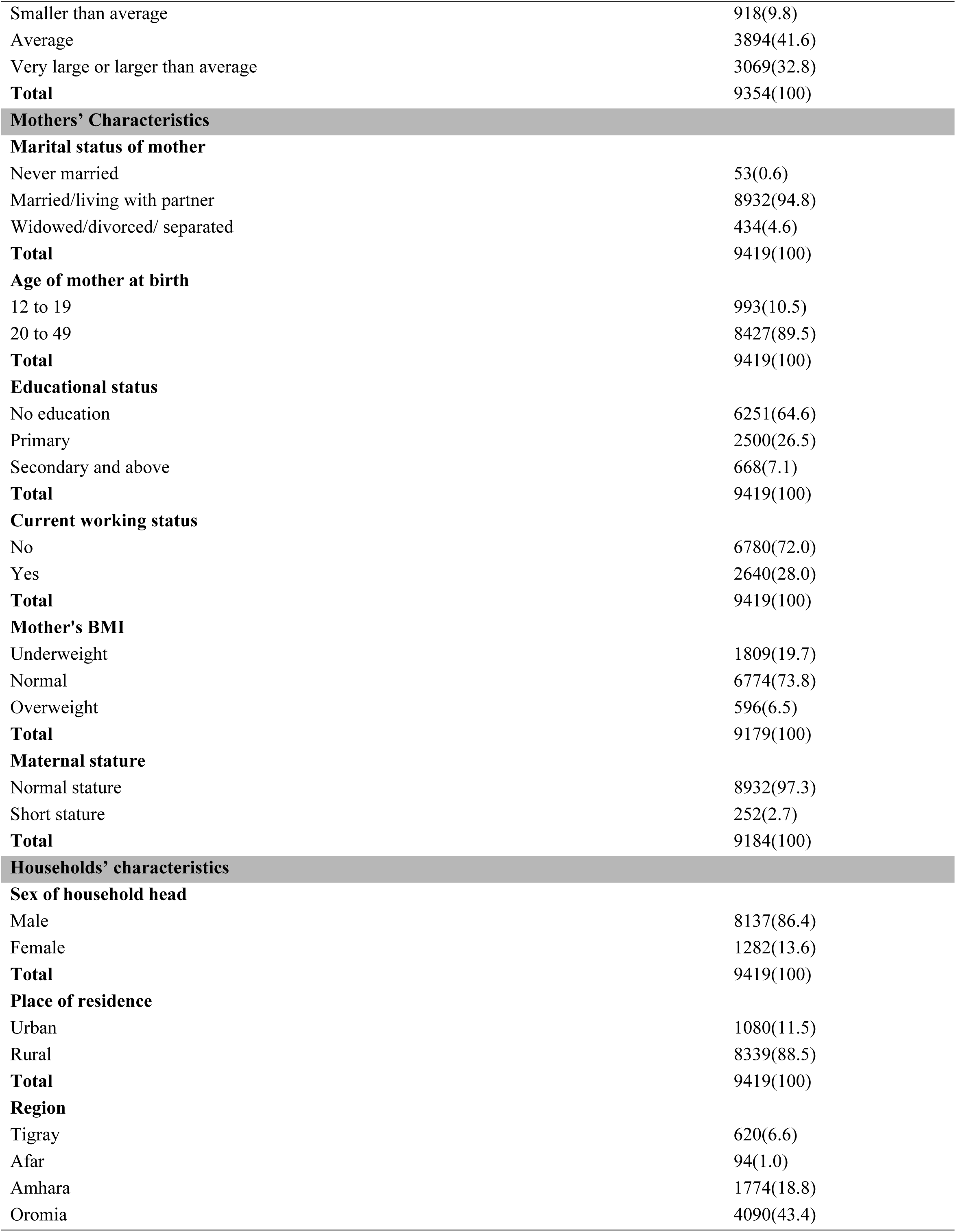
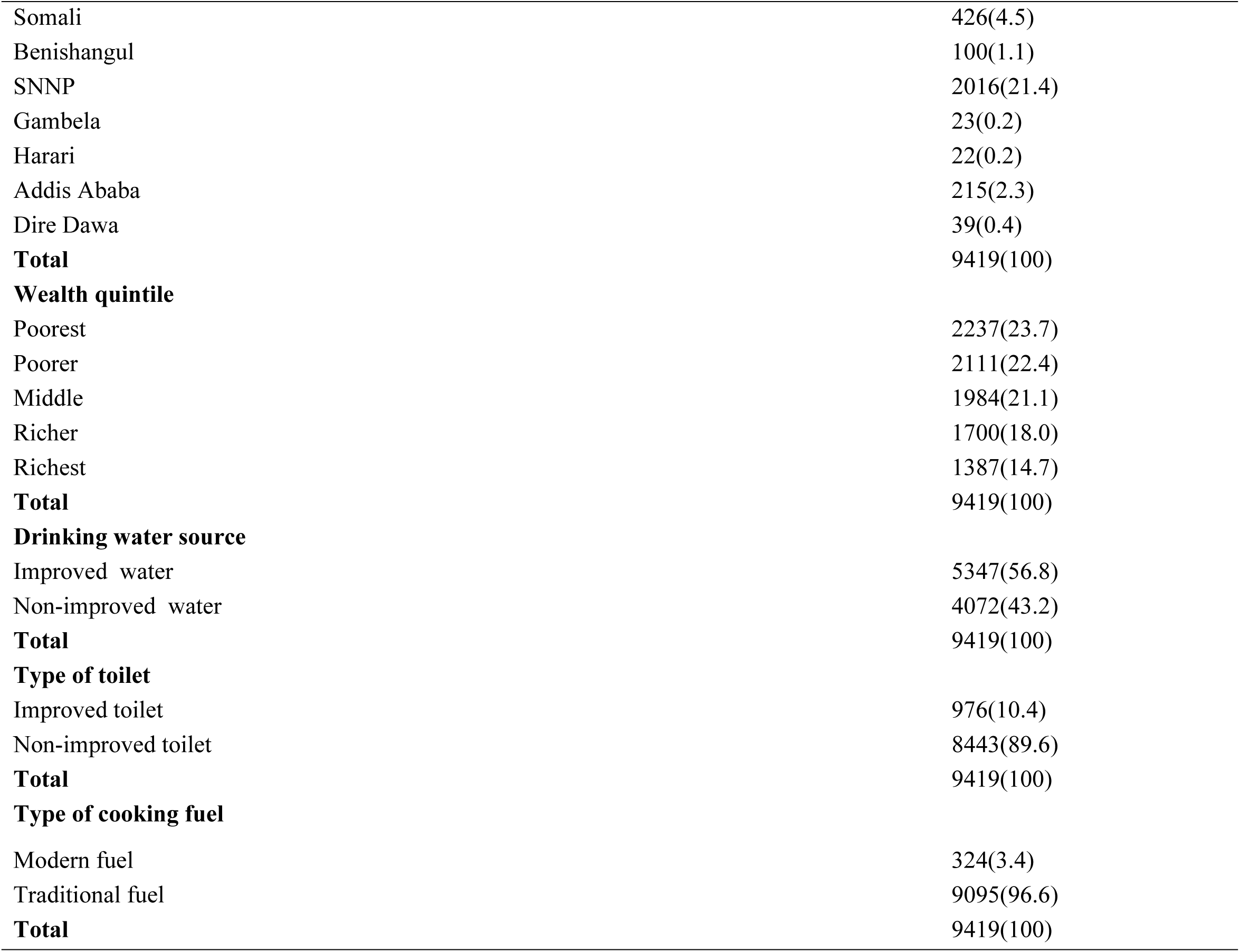
Percent distribution of sampled children age 0 to 59 months by child’s, mothers’, household, and environment characteristics, Ethiopia DHS 2016

| Variables | N (%) |
| --- | --- |
| **Child's Characteristics** |  |
| **Child age** |  |
| less than 6 months | 703(7.5) |
| 6 to 11 months | 1018(10.8) |
| 12 to 17 months | 1105(11.7) |
| 18 to 23 | 831(8.8) |
| 24 to 59 | 5762(61) |
| **Total** | 9419(100) |
| **Sex of child** |  |
| Male | 4832(51.3) |
| Female | 4587(48.7) |
| **Total** | 9419(100) |
| **Birth order** |  |
| First | 1656(17.6) |
| 2 to 4 | 4066(43.2) |
| Greater than or equal to 5 | 3697(39.3) |
| **Total** | 9419(100) |
| **Perceived weight at birth** |  |
| Very small | 1474(15.8) |
| Smaller than average | 918(9.8) |
| Average | 3894(41.6) |
| Very large or larger than average | 3069(32.8) |
| **Total** | 9354(100) |
| **Marital status of mother** |  |
| Never married | 53(0.6) |
| Married/living with partner | 8932(94.8) |
| Widowed/divorced/ separated | 434(4.6) |
| **Total** | 9419(100) |
| **Age of mother at birth** |  |
| 12 to 19 | 993(10.5) |
| 20 to 49 | 8427(89.5) |
| **Total** | 9419(100) |
| **Educational status** |  |
| No education | 6251(64.6) |
| Primary | 2500(26.5) |
| Secondary and above | 668(7.1) |
| **Total** | 9419(100) |
| **Current working status** |  |
| No | 6780(72.0) |
| Yes | 2640(28.0) |
| **Total** | 9419(100) |
| **Mother's BMI** |  |
| Underweight | 1809(19.7) |
| Normal | 6774(73.8) |
| Overweight | 596(6.5) |
| **Total** | 9179(100) |
| **Maternal stature** |  |
| Normal stature | 8932(97.3) |
| Short stature | 252(2.7) |
| **Total** | 9184(100) |
| **Sex of household head** |  |
| Male | 8137(86.4) |
| Female | 1282(13.6) |
| **Total** | 9419(100) |
| **Place of residence** |  |
| Urban | 1080(11.5) |
| Rural | 8339(88.5) |
| **Total** | 9419(100) |
| **Region** |  |
| Tigray | 620(6.6) |
| Afar | 94(1.0) |
| Amhara | 1774(18.8) |
| Oromia | 4090(43.4) |
| Somali | 426(4.5) |
| Benishangul | 100(1.1) |
| SNNP | 2016(21.4) |
| Gambela | 23(0.2) |
| Harari | 22(0.2) |
| Addis Ababa | 215(2.3) |
| Dire Dawa | 39(0.4) |
| **Total** | 9419(100) |
| **Wealth quintile** |  |
| Poorest | 2237(23.7) |
| Poorer | 2111(22.4) |
| Middle | 1984(21.1) |
| Richer | 1700(18.0) |
| Richest | 1387(14.7) |
| **Total** | 9419(100) |
| **Drinking water source** |  |
| Improved water | 5347(56.8) |
| Non-improved water | 4072(43.2) |
| **Total** | 9419(100) |
| **Type of toilet** |  |
| Improved toilet | 976(10.4) |
| Non-improved toilet | 8443(89.6) |
| **Total** | 9419(100) |
| **Type of cooking fuel** |  |
| Modern fuel | 324(3.4) |
| Traditional fuel | 9095(96.6) |
| **Total** | 9419(100) |

### Bivariate associations of stunting and wasting by background characteristics

Table 2 indicates results from the bivariate analyses. Child age, Sex, birth order, perceived weight, mothers’ education, body mass index, place of residence, region, wealth quintile, toilet and fuel type were significantly associated with stunting. Variables associated with wasting include child age, perceived weight, body mass index, region, wealth quintile, and fuel type.

**Table 2.** Bivariate analysis stunting and wasting by background characteristics

| Independent Variables | Outcome variables |  |  |  |
| --- | --- | --- | --- | --- |
|  | Stunting<br>Percentage | 95% CI | Wasting<br>Percentage | 95% CI |
| **Children's characteristics** |  | **P=0.000** |  | **P=0.000** |
| **Child Age** |  |  |  |  |
| Less than 6 months | 13.6 | [10.0,18.3] | 15.5 | [11.5,20.5] |
| 6 to 11 months | 17.2 | [13.4,21.7] | 12.9 | [10.1,16.4] |
| 12 - 17 | 31.7 | [27.7,36.1] | 15.4 | [12.3,19] |
| 18 - 23 | 48.6 | [43.2,54.1] | 11.3 | [7.8,16.0] |
| 24 - 59 | 45.5 | [42.9,48.1] | 7.5 | [6.5,8.6] |
| **Total N=** | **3373** |  | **875** |  |
| **Sex of Child** |  | **P=0.001** |  | **P=0.760** |
| Male | 41.2 | [38.7,43.8] | 10.1 | [8.8,11.6] |
| Female | 35.7 | [33.4,38.2] | 9.8 | [8.4,11.4] |
| **Total N=** | **3372** |  | **874** |  |
| **Birth Order** |  | **P=0.039** |  | **P=0.170** |
| First | 36.1 | [31.8,40.7] | 8.3 | [6.4,10.7] |
| 2 to 4 | 37.0 | [34.4,39.7] | 9.7 | [8.2,11.5] |
| Greater than or equal to 5 | 41.3 | [38.5,44.3] | 11.0 | [9.5,12.7] |
| **Total N=** | **3372** |  | **875** |  |
| **Perceived Weight at Birth** |  | **P=0.000** |  | **P=0.002** |
| Very small | 46.2 | [41.5,51] | 13.4 | [10.9,16.5] |
| Smaller than average | 43.6 | [38.7,48.6] | 12.0 | [9.2,15.6] |
| Average | 37.6 | [34.9,40.3] | 9.6 | [8.2,11.2] |
| Very large or larger than average | 34.6 | [31.8,37.5] | 8.2 | [6.8,9.9] |
| **Total N=** | **3355** |  | **870** |  |
| **Mothers' Characteristics** |  |  |  |  |
| **Marital Status of Mother** |  | **P=0.771** |  | **P=0.531** |
| Never married | 44.5 | [26.8,63.8] | 5.2 | [0.8,28.1] |
| Married/living with partner | 38.4 | [36.5,40.4] | 9.9 | [8.9,11] |
| Widowed/divorced/ separated | 40.6 | [33.2,48.4] | 12.0 | [7.8,17.9] |
| **Total N=** | **3373** |  | **875** |  |
| **Age of Mother at birth** |  | **P=0.111** |  | **P=0.478** |
| 12 to 19 | 42.5 | [37.2,48] | 9.0 | [6.7,12] |
| 20 to 49 | 38.1 | [36.1,40.5] | 10.1 | [9.0,12.2] |
| **Total N=** | **3373** |  | **875** |  |
| **Educational Status** |  | **P=0.000** |  | **P=0.051** |
| No education | 41.7 | [39.5,44] | 10.8 | [9.5,12.2] |
| Primary | 35.6 | [32.5,38.8] | 8.7 | [7.1,10.7] |
| Secondary and above | 19.5 | [15.3,24.4] | 7.2 | [4.8,10.7] |
| **Total N=** | **3373** |  | **875** |  |
| **Working status of mother** |  | **P=0.947** |  | **P=0.410** |
| No | 38.5 | [36.3,40.8] | 9.7 | [8.5,11.0] |
| Yes | 38.6 | [35.7,41.7] | 10.7 | [8.8,12.9] |
| **Total N=** | **3373** |  | **875** |  |
| **Mother's BMI** |  | **P=0.000** |  | **P=0.000** |
| Underweight | 41.9 | [38.5,45.5] | 13.5 | [11.3,16] |
| Normal | 38.9 | [36.5,41.3] | 9.5 | [8.4,16] |
| Overweight | 24.4 | [19.6,29.9] | 4.6 | [2.4,8.8] |
| **Total N=** | **3342** |  | **867** |  |
| **Maternal stature** |  | **P=0.000** |  | **P=0.396** |
| Normal stature | 38.1 | [36.1,40.1] | 9.9 | [8.8,11.1] |
| Short stature | 54.8 | [45.5,63.7] | 12.6 | [7.4,20.7] |
| **Total N=** | **3342** |  | **867** |  |
| **Households' Characteristics** |  |  |  |  |
| **Sex of Household Head** |  | **P=0.2196** |  | **P=0.287** |
| Male | 38.4 | [36.4,40.6] | 10.1 | [9.0,11.4] |
| Female | 39.2 | [35.7,42.8] | 8.8 | [7.0,11.1] |
| **Total N=** | **3372** |  | **875** |  |
| **Place of Residence** |  | **P=0.000** |  | **P=0.509** |
| Urban | 26.0 | [21.5,31.1] | 9.1 | [7.0,11.8] |
| Rural | 40.1 | [38.1,42.2] | 10.1 | [9.0,11.3] |
| **Total N=** | **3373** |  | **875** |  |
| **Region** |  | **P=0.000** |  | **P=0.000** |
| Tigray | 39.5 | [35.8,43.4] | 11.5 | [9.5,13.8] |
| Afar | 40.9 | [36.5,45.5] | 17.1 | [13.4,21.6] |
| Amhara | 47.6 | [43.8,51.4] | 10.0 | [8.0,12.5] |
| Oromia | 36.1 | [32.8,39.5] | 10.5 | [8.8,12.6] |
| Somali | 26.5 | [22.8,30.6] | 22.2 | [17.5,27.8] |
| Benishangul | 42.9 | [38.2,47.6] | 10.5 | [7.8,14.0] |
| SNNP | 39.6 | [35.7,43.6] | 6.2 | [4.8,8.0] |
| Gambela | 24.1 | [20.3,28.4] | 13.9 | [10.4,18.4] |
| Harari | 32.1 | [27.7,36.8] | 10.7 | [7.4,15.2] |
| Addis Ababa | 14.9 | [11.1,19.7] | 2.4 | [1.3,4.6] |
| Dire Dawa | 40.9 | [34.5,47.6] | 10.2 | [7.1,14.4] |
| **Total N=** | **3374** |  | **874** |  |
| **Wealth quintile** |  | **P=0.000** |  | **P=0.001** |
| Poorest | 45.6 | [42.4,48.8] | 13.8 | [11.2,16.8] |
| Poorer | 42.5 | [38.7,46.5] | 9.6 | [7.7,12.0] |
| Middle | 38.6 | [34,43.4] | 10.4 | [8.2,13.1] |
| Richer | 34.8 | [31,38.7] | 6.8 | [5.1,9.0] |
| Richest | 25.7 | [22.1,29.7] | 7.7 | [5.7,10.2] |
| **Total N=** | **3373** |  | **874** |  |
| **Drinking Water source** |  | **P=0.048** |  | **P=0.098** |
| Improved water | 37.0 | [34.6,39.5] | 9.2 | [8.0,10.5] |
| Non-Improved water | 40.5 | [37.8,43.3] | 11.0 | [9.3,12.8] |
| **Total N=** | **3372** |  | **875** |  |
| **Toilet Facilities** |  | **P=0.000** |  | **P=0.222** |
| Improved toilet | 26.8 | [22.4,31.7] | 8.5 | [6.5,11.1] |
| Non-improved toilet | 39.8 | [37.8,42] | 10.0 | [9.0,11.3] |
| **Total N=** | **3373** |  | **874** |  |
| **Type of Cooking Fuel** |  | **P=0.000** |  | **P=0.049** |
| Modern fuel | 21.4 | [14.9,29.9] | 5.2 | [2.6,10.2] |
| Traditional fuel | 39.1 | [37.2,41.1] | 10.1 | [9.1,11.3] |
| **Total N=** | **3373** |  | **875** |  |

### Factors Associated with Stunting

Table 3 shows the results of the adjusted multiple logistic regressions for stunning and wasting. Child age has a significant association with stunting. Older Children were more likely to be stunted compared to less than age 6months. The odds were higher in the age group, 24-59(AOR=6.59; CI 5.00-8.68), 18-23(AOR=7.81; CI 5.73-10.65), and age 12-17(AOR=3.50; CI 2.59-1.86). Female children were less likely to be stunted than male children (AOR=0.82; CI 0.74-0.90). Child at birth with very large or larger than average in their perceived weight have less likely to be stunted compared to very small children at birth (AOR=0.54; CI 0.47-0.63). Very large or larger than average (AOR=0.54; CI 0.47-0.63), average (AOR=0.68; CI 0.59-0.78), smaller than average (AOR=0.82; CI 0.67-0.99). Mothers with educational status of secondary and above were less likely to have stunted children (AOR=0.66; CI 0.52-0.84) than mothers with no education.

**Table 3.** Prevalence of stunting and wasting among children age 0 to 59 months by child’s mothers’, households and environment characteristics

| Independent Variables |  | Outcome Variables |  |
| --- | --- | --- | --- |
|  |  | Stunting<br>AOR (95%CI) | Wasting<br>AOR (95%CI) |
| **Children's characteristics** |  |  |  |
| Child Age | Less than 6 months | 1 | 1 |
|  | 6 to 11 months | 1.34(0.97 - 1.86) | 1.10(0.82 - 1.50) |
|  | 12 to 17 | 3.50*** (2.59 - 4.74) | 1.13(0.84 - 1.52) |
|  | 18 - 23 | 7.81*** (5.73 - 10.65) | 0.87(0.62 - 1.21) |
|  | 24 - 59 | 6.59*** (5.00 - 8.68) | 0.56*** (0.43 - 0.73) |
| Sex of Child | Male | 1 | 1 |
|  | Female | 0.82*** (0.74 - 0.90) | 0.77*** (0.67 - 0.88) |
| Birth Order | First | 1 | 1 |
|  | 2 to 4 | 1.02(0.87 - 1.20) | 1.02(0.80 - 1.29) |
|  | Greater than or equal to 5 | 1.16(0.97 - 1.39) | 1.16(0.90 - 1.50) |
| Perceived weight at Birth | Very small | 1 | 1 |
|  | Smaller than average | 0.82*(0.67 - 0.99) | 1.07(0.83 - 1.37) |
|  | Average | 0.68*** (0.59 - 0.78) | 0.71*** (0.59 - 0.86) |
|  | Very large or larger than average | 0.54*** (0.47 - 0.63) | 0.60*** (0.49 - 0.74) |
| **Mothers' Characteristics** |  |  |  |
| Marital Status of Mother | Never in union | 1 | 1 |
|  | Married/living with partner |  | 2.16(0.51 - 9.14) |
|  | Widowed/divorced/ separated | 0.64(0.32 - 1.26) | 2.48(0.57 - 10.76) |
| Age of Mother | 12 to 19 | 1 | 1 |
|  | 20 to 49 | 0.97(0.81 - 1.16) | 0.96(0.74 - 1.25) |
| Educational Status | No education |  |  |
|  | Primary | 0.97(0.86 - 1.10) | 0.92(0.76 - 1.11) |
|  | Secondary and above | 0.66*** (0.52 - 0.84) | 0.80(0.57 - 1.13) |
| Current working status | No | 1 | 1 |
|  | Yes | 1.02(0.91 - 1.14) | 1.08(0.91 - 1.28) |
| Mother's BMI | underweight |  |  |
|  | Normal | 0.84** (0.74 - 0.94) | 0.67*** (0.57 - 0.78) |
|  | Overweight | 0.57*** (0.45 - 0.71) | 0.39*** (0.28 - 0.55) |
| Maternal stature | Normal stature | 1 | 1 |
|  | Short stature | 2.03*** (1.44 - 2.86) | 1.08(0.65 - 1.80) |
| **Households' Characteristics** |  |  |  |
| Sex of Household Head | Male | 1 | 1 |
|  | Female | 1.07(0.94 - 1.23) | 0.97(0.81 - 1.17) |
| Type of Place of Residence | Urban |  |  |
|  | Rural | 0.86(0.69 - 1.07) | 0.61(0.64 - 1.10) |
| Region | Tigray | 1 | 1 |
|  | Afar | 0.87(0.69 - 1.10) | 1.33(0.97 - 1.81) |
|  | Amhara | 1.27*(1.03 - 1.58) | 0.87(0.62 - 1.20) |
|  | Oromia | 0.83(0.68 - 1.01) | 1.00(0.75 - 1.34) |
|  | Somali | 0.52*** (0.42 - 0.65) | 2.02*** (1.51 - 2.68) |
|  | Benishangul | 1.13(0.90 - 1.43) | 0.97(0.69 - 1.37) |
|  | SNNP | 0.99(0.80 - 1.21) | 0.60** (0.43 - 0.85) |
|  | Gambela | 0.51*** (0.39 - 0.66) | 1.19(0.84 - 1.67) |
|  | Harari | 0.87(0.67 - 1.15) | 1.20(0.81 - 1.76) |
|  | Addis Ababa | 0.64* (0.43 - 0.94) | 0.35** (0.17 - 0.76) |
|  | Dire Dawa | 1.09(0.83 - 1.43) | 0.83(0.55 - 1.27) |
| Wealth quintile | Poorest | 1 | 1 |
|  | Poorer | 1.01(0.86 - 1.17) | 0.88(0.71 - 1.10) |
|  | Middle | 0.77** (0.65 - 0.91) | 0.92(0.72 - 1.16) |
|  | Richer | 0.69*** (0.58 - 0.82) | 0.66** (0.50 - 0.87) |
|  | Richest | 0.62*** (0.49 - 0.79) | 0.58** (0.40 - 0.83) |
| Drinking water source | Improved water | 1 | 1 |
|  | Non-improved water | 0.91(0.81 - 1.01) | 1.00(0.86 - 1.17) |
| Type of toilet Facilities | Improved toilet | 1 | 1 |
|  | Non-improved toilet | 1.28** (1.07 - 1.53) | 1.06(0.83 - 1.36) |
| Type of Cooking Fuel | Modern fuel | 1 | 1 |
|  | Traditional fuel | 1.66** (1.18 - 2.32) | 1.28(0.74 - 2.21) |

\*\*\* p<0.001, \*\* p<0.01, \* p<0.05 AOR, adjusted odds ratio

Children born from overweight mothers (AOR=0.57; CI 0.5-0.71) and normal mothers in their bod mass index were less likely to be stunted (AOR=0.84; CI 0.74-0.94) than born from underweight mothers. Children from short stature mothers were more likely to be stunted (AOR=2.03; CI 1.44-2.86) than children born from not short stature mothers. There is significant regional variation in stunting among under five children in Ethiopia. Children from Amhara region were more likely to be stunted (AOR=1.27; CI 1.03-1.58) than Tigray region. Children from Addis Abeba (AOR=0.64; CI 0.43-0.94), Gambella (AOR=0.51; CI 0.39-0.66), and Somali (AOR=0.52; CI 0.42-0.65) were less likely to be stunted than Tigray region of Ethiopia. Children from the richest (AOR=0.62; CI 0.49-0.79), richer (AOR=0.69; CI 0.58-0.82), and middle income households (AOR=0.77; CI 0.65-0.91) were less likely to be stunted than children from poorer and poorest households. Children from non-improved toilet user households were more likely to be stunted (AOR=1.28; CI 1.07-1.53) than born from households having improved toilet users. Children from traditional fuel user households were more likely to be stunted (AOR=1.66; CI 1.18-2.32).

### Factors Associated with Wasting

Children from 24-59 months of age group were less likely to be wasted (AOR=0.56; CI 0.43-0.73) than 0-6 months. Children with larger than average and average in their mothers’ perceived weight were less likely to be wasted with the odds of (AOR=0.60; CI 0.49-0.74) and (AOR=0.71; CI 0.59-0.86) respectively. Female children were less likely to be wasted than male (AOR=0.77; CI 0.67-0.88). Overweight mothers (AOR=0.39; CI 0.28-0.55) and normal (AOR=0.67; CI 0.57-0.78) in their body mass index have less wasting than children from underweight mothers. Children born from rural residents were less likely to be wasted than children born from urban resident households (AOR=0.61; CI 0.45-0.83). Households reside in Addis Abeba (AOR=0.35; CI 0.17-0.76) and SNNP (AOR=0.60; CI 0.43-0.85) were less likely to have wasted children. Households from Somali region were more likely to have wasted children (AOR=2.02; CI 1.51-2.68).

## DISCUSSION

This study assessed the determinants of nutritional status among children under five in Ethiopia. Study results show child age, sex, perceived birth weight, mothers’ educational status, body mass index, maternal stature, region, wealth index, toilet and cooking fuel types had significant association with stunting. Similarly child age, sex, perceived birth weight, body mass index, residence and region have also significant associations with wasting.

This study shows, the risk of stunting increased with age of child. Similarly previous studies in Ethiopia such as Genebo, Girma (15), Asfaw and Giotom (16)and Yimer (17) had similar results. The reason might be greater energy needs as child age increased. Besides, this could be due to stunting is a chronic malnutrition that can be manifested after long-term nutritional deficiency and wasting reflects acute undernutrition. As child age increase there is probability to finish vaccine which reduce exposure to disease. For example Mekonnen, Jones (18) found children vaccinated against measles are less likely to be wasted. In many developing countries including Ethiopia, complementary feeding is a challenge to children age 6 to 23months (19). Studies in other developing countries by Caulfield, Huffman (20), Mora and Nestel (21)and Abeshu, Lelisa (19) found that complementary foods fed to infants in the second six months of age and beyond are often inadequate in energy density.

Female children were less likely to be stunted and wasted than boys. This finding is consistent with a meta-analysis in Sub-Saharan Africa (22), a study in the Northern Ethiopia (23) and Myanmar(24). However, this study is in contrary to studies in Tanzania (25),Pakistan (26), India (27), and Kenya (28) that found girls had a higher prevalence in stunting than boys. Garenne (29), Wamani, Åstrøm (22), Bork and Diallo (30) found the sex differences in nutritional status might be due to biological differences in morbidity between boys and girls in their early life. This may also apply to our findings on wasting. A review of complementary feeding reported that, boys have higher birth weights than girls and grow faster during infancy, resulting in greater energy needs(30). Furthermore another study on Guatemalan infants reported that women perceived male children to be hungrier and less likely to be satisfied by breastfeeding alone (31), placing them at increased risk of infections.

Regarding perceived weight of child at birth, the risks of stunting and wasting were higher among children reported low birth weight. The findings are consistent with previous studies in Zambia (32), Bangladesh (33-35), India (36), Malaysia(37),and Indonesia(38) and may be due to the poor nutrition of mothers during pregnancy. This finding is supported by our results of the association between mother’s body mass index and stunting and wasting in children. Ramakrishnan (39)reported that children born from overweight mothers are less likely to be stunted and wasted due to adequacy of protein and energy intakes of mothers during pregnancy(39). Similar studies in Nepal (40) also shows as mothers’ body mass index increase, there was a reduction in child’s stunting, and wasting. Our study found that children born from mothers with short stature are more likely to be stunted. This result was in line with previous studies from (41) (42). This is due to women who were stunted during their childhood are more likely to give birth to stunted children(43).

Higher Educational status of mothers was positively associated with stunting. This is in line with previous related studies in Philippines(44), Libya (45), Ethiopia (17), Uganda (46), Mozambique (47), and Ghana (42). The reason was stated in different literatures; educated women are better informed about optimal child care practices (48), have better practices in terms of hygiene (49, 50) feeding (51),and childcare during illness (51-53), have a greater ability to use the health system (54), are more empowered to make decisions (49). However, the majority (65%) of sampled households’ did not attend formal education as a result knowledge of mothers on child care may not be enough.

This study finding shows there is regional variation in stunting and wasting. Children from Addis Ababa were less likely to be stunted and wasted. While children in Gambella and SNNP were less likely to be stunted and wasted respectively. Interestingly, children in the Somali regions were less likely to be stunted, but more likely to be wasted. This could be due to wasting is characterized by acute malnutrition that can be caused by temporary increased food insecurity from extreme weather events, drought, and shifts in agricultural practices (55, 56), and studies from Ethiopia by Pelletier, Deneke (57) shows association between malnutrition was varies across regions due to variation in cultivated area. In the DHS survey households’ wealth were usually measured by increments in household material standards by calculating wealth index. Previous studies in Nepal (58) and other Asian countries (59) found that household asset accumulation is an important predictors of nutritional improvement in most countries.

Our study found that children from non-improved toilet and traditional fuel user households were more likely to be stunted. Environmental factors such as unimproved water, unimproved sanitation, and biomass fuel are the second largest global attributable burden as cause of child stunting (60). A study from 137 developing countries reported that 7.2 million cases of stunting were attributable to unimproved sanitation (60). In Ethiopia more than half (56%) of rural households use unimproved toilet facilities (10). One in three households in Ethiopia have no toilet facility (39%) in rural areas and 7% in urban areas (10). Related studies (61-64) in Sri Lanka revealed that malnutrition among children is still a major health problem associated with poor sanitation and personal hygiene. However, study conducted in rural Zimbabwe by Humphrey, Mduduzi N N Mbuya (65) found improved water, sanitation and hygiene interventions implemented in rural areas in low-income countries were unlikely to reduce stunting more than implementation of improved infant and child feeding alone.

This implies integration of improved infant diets with improved water, sanitation and hygiene is a logical approach to improve nutritional status of under five children. The majority (93%) of Ethiopian households use traditional fuels such as fuel wood, charcoal, and dung cake to meet their daily needs and most households (59%) cook food outside(10). Ramakrishnan (39) found that indoor air pollution and smoking are causes of low birth weight and could possibly impact stunting. While another study in India reported associations between air pollution and stunting(66). Children living in households using biofuel were more likely to be stunted compared to children living in households using cleaner fuels(66).

## STRENGTHS AND LIMITATIONS

The strength of this study were; it used national representative data which allow to generalize the results. It also applied all the DHS data principles like weighting. Regarding the limitations, the cross-sectional data cannot examine causation and seasonal variation of nutritional outcomes. The other limitation is that during DHS data collection, measurements of height by laying down under 2 children may considered as source of human error.

## CONCLUSION AND POLICY IMPLICATIONS

Despite efforts made by the Ethiopian government and improvements in reducing malnutrition, rates of stunting and wasting remain high. The consequences of child malnutrition are grave and include poor performance in school and low adult economic productivity. The study found child, maternal, household and environmental characteristics were significantly associated with stunting and wasting among of children under five. The findings imply that a multi-sectorial and multidimensional approach is important to address malnutrition in Ethiopia. The education sector should promote maternal education and policies to reduce cultural and gender barriers. The health sector should provide health education to families to change behaviors of child care givers towards child care and infant feeding practices. In addition, the health and human services sectors can promote use of improved toilets. To reduce stunting due to traditional fuel usage in the households, the energy sector should help households to access clean energy. Further research is suggested on the impact of seasonal food insecurity and climatic events on stunting and wasting and research to examine the associated factors on nutritional status among children over time using decomposition analysis.

## Acknowledgments

Foremost, we would like to thank USAID for the DHS data analysis Training through the 2019 DHS Fellows program implemented by ICF. We thanks to our training facilitators Dr. Shireen Assaf and Dr. Wenjuan Wang. Their scholarly guidance and encouragement was the source of energy for our research. We would like to express our special thanks to our co-facilitators, Mr. Gedefaw Abeje Fekade (Assistant Professor), and Dr. Kyaw Swa Mya for their invaluable assistance to improve our research work during the training. We especially appreciate the support of Dr. Sorrel Namaste for her advice and sharing valuable materials at the inception that supports our inclusion criteria during sampling procedure. We also appreciate Dr.Rukundo Benedict and Yodit Bekele for their valuable comments and their time spent in sharing related materials to improve our work.

